# Pressure and ultrasound activate mechanosensitive TRAAK K^+^ channels through increased membrane tension

**DOI:** 10.1101/2023.01.11.523644

**Authors:** Ben Sorum, Trevor Docter, Vincent Panico, Robert A. Rietmeijer, Stephen G. Brohawn

## Abstract

TRAAK is a mechanosensitive two-pore domain K^+^ (K2P) channel found in nodes of Ranvier within myelinated axons. It displays low leak activity at rest and is activated up to one hundred-fold by increased membrane tension. Structural and functional studies have led to physical models for channel gating and mechanosensitivity, but no quantitative analysis of channel activation by tension has been reported. Here, we use simultaneous patch-clamp recording and fluorescent imaging to determine the tension response characteristics of TRAAK. TRAAK shows high sensitivity and a broad response to tension spanning nearly the entire physiologically relevant tension range. This graded response profile distinguishes TRAAK from similarly low-threshold mechanosensitive channels Piezo1 and MscS, which activate in a step-like fashion over a narrow tension range. We further use patch imaging to show that ultrasonic activation of TRAAK and MscS is due to increased membrane tension. Together, these results provide mechanistic insight into TRAAK tension gating, a framework for exploring the role of mechanosensitive K^+^ channels at nodes of Ranvier, and biophysical context for developing ultrasound as a mechanical stimulation technique for neuromodulation.

## Introduction

Mechanosensitive ion channels are opened by mechanical force to rapidly transduce physical stimuli into cellular electrical signals^1-3^. Their activity underlies a wide range of physiological processes, from the classic senses of touch and audition to proprioception, blood pressure regulation, digestion, osmolarity control, and cell growth. Known mechanosensitive channels are diverse. They belong to evolutionarily distinct families and exhibit varying ion selectivity, kinetics, conductance, force-gating mechanisms, and sensitivity to mechanical gating stimuli^1,2^. Though characterization of these properties is essential to understanding how forces are sensed and encoded, quantification of mechanosensitive channel responses to gating forces remains limited.

TRAAK is a mechanosensitive member of the two-pore domain (K2P) K^+^ ion channel family^15^ and is localized to nodes of Ranvier, the small gaps between myelinated regions of axons where the action potential is regenerated during saltatory conduction^19–21^. In contrast to canonical models for the molecular basis of the action potential like the squid giant axon, mammalian axons can completely lack voltage-gated K^+^ channels at nodes^22^. Instead, TRAAK and the related mechanosensitive K2P TREK1 contribute to the large K^+^ conductance at mammalian nodes that sets the resting potential, repolarizes the membrane in an action potential, and maintains voltage-gated Na^+^ channel availability to facilitate high-frequency spiking^19–21^. Disruption of TRAAK activity has physiological implications: knockout mice show mechanical allodynia and hyperalgesia while gain-of-function mutations in humans cause the severe neurodevelopmental disorder FHEIG (facial dysmorphism, hypertrichosis, epilepsy, intellectual disability/developmental delay, and gingival overgrowth)^23,24^. TRAAK displays low basal leak activity under resting conditions but is activated up to ∼100-fold by increased membrane tension^16,17^. Leak and mechanically gated activity arise from physically distinct open states^16^. At low tension, TRAAK is predominantly closed due to lipid block of the pore^18^. Delipidation produces leak activity, while mechanically gated activity involves conformational changes—most notably the upward movement of transmembrane helix 4 towards the extracellular solution—that seal laterally facing membrane openings to prevent lipid block^16,18^. These conformational changes increase channel cross-sectional area and cylindricity, shape changes that are energetically favored by increased tension^18^. Whether and how mechanical gating of TRAAK contributes to its role at nodes remains unknown. A quantitative understanding of TRAAK response to tension is an important step towards clarifying potential models for channel function at nodes and understanding the mechanistic basis for its mechanosensitivity.

Membrane tension is the gating stimulus for many mechanosensitive ion channels^3,4^. Still, tension is not measured in typical assays of channel activity. Instead, experimentally measured stimulation parameters are indirect: probe displacement during cell poking, substrate elongation or pillar displacement during cell stretching, osmolarity during cell swelling, and pressure during patched membrane stretching. These approaches are thought to increase membrane tension in addition to causing other assay-specific effects. Moreover, relating tension to the most readily measured parameters is non-trivial. One solution is to simultaneously electrophysiologically record and image patched membranes during pressure-induced activation of mechanosensitive channels^5–14^. Membrane tension (*T*) can then be calculated according to the Young-Laplace equation (*T* = ΔPr/2) using values of applied pressure (ΔP) and measured membrane radius of curvature (r). This approach has been successfully utilized in studies of Piezo1, MscS, and MscL^5–14^.

An emerging approach for activating mechanosensitive channels is applying low-power and low-frequency ultrasound stimulation^25,26^. Activation of mechanosensitive ion channels may underlie ultrasound’s neuromodulatory properties^17,27–29^, which were first identified nearly a century ago and have been widely reported across the nervous system since. Ultrasound is advantageous for neuromodulation and manipulating ion channel activity as it can be delivered and focused non-invasively through tissue and bone^25,26^. We previously showed that ultrasound activates TRAAK in patches from cells and proteoliposomes and specifically promotes the same mechanically gated open state as pressure stimulation^17^. This suggests ultrasound could activate mechanosensitive channels by increasing membrane tension rather than through alternative means of energy transfer.

Here, we develop and implement a simultaneous patch imaging and recording assay to quantify the tension response of TRAAK and show that ultrasound stimulation, like pressure stimulation, stretches patched membranes, consistent with tension-mediated activation of mechanosensitive channels.

## Results

We expressed human TRAAK in *Xenopus laevis* oocytes and recorded currents across excised (inside-out) patches in response to mechanical stimulation generated by pressure steps to the base of the patch pipette (Figure 1A). Currents were recorded at 0 mV in a 10-fold gradient of K^+^ across the membrane. Patches contained thousands of channels given the TRAAK single channel current of ∼1-2 pA under these conditions^16^. We normalized the peak current from each pressure step to the maximum current recorded from that patch in response to mechanical stimulation (I/I_max_) to compile data across patches (data from a subset of recordings are displayed in Fig. 1B). Patches were only analyzed if a saturating current response was observed. TRAAK activity was low under resting conditions and activated by pressure up to an average of 41.6 ± 21.6-fold (mean ± SEM, n = 18 patches), similar to prior reports^16,17^. Data were fit to a Boltzmann with a P_50_ = -6.2 ± 0.9 mmHg and 10 to 90% range of -1.7 to -10.7 mmHg (mean ± SEM, n = 18 patches). As a comparison, we also recorded currents from the well-characterized *E. coli* mechanosensitive channel MscS (Fig. 1C)^30^. MscS was activated by similar negative pressures to TRAAK (P_50_ = -6.5 ± 0.8 mmHg, 10 to 90% range -1.8 to -10.7 mmHg) up to an average of 14.6 ± 4.0-fold (mean ± SEM, n= 10 patches) (data from a subset of recordings are displayed in Fig. 1D). Patches contained tens of MscS channels given a single channel current of ∼12 pA at -60 mV under these conditions^31^. Both TRAAK and MscS displayed substantial patch-to-patch variability in pressure response (Fig. 1B, D, fit R^2^ = 0.49 and 0.64, respectively). This is consistent with membrane tension, rather than pressure, being the stimulus that promotes channel opening^11^.

**Figure 1.**
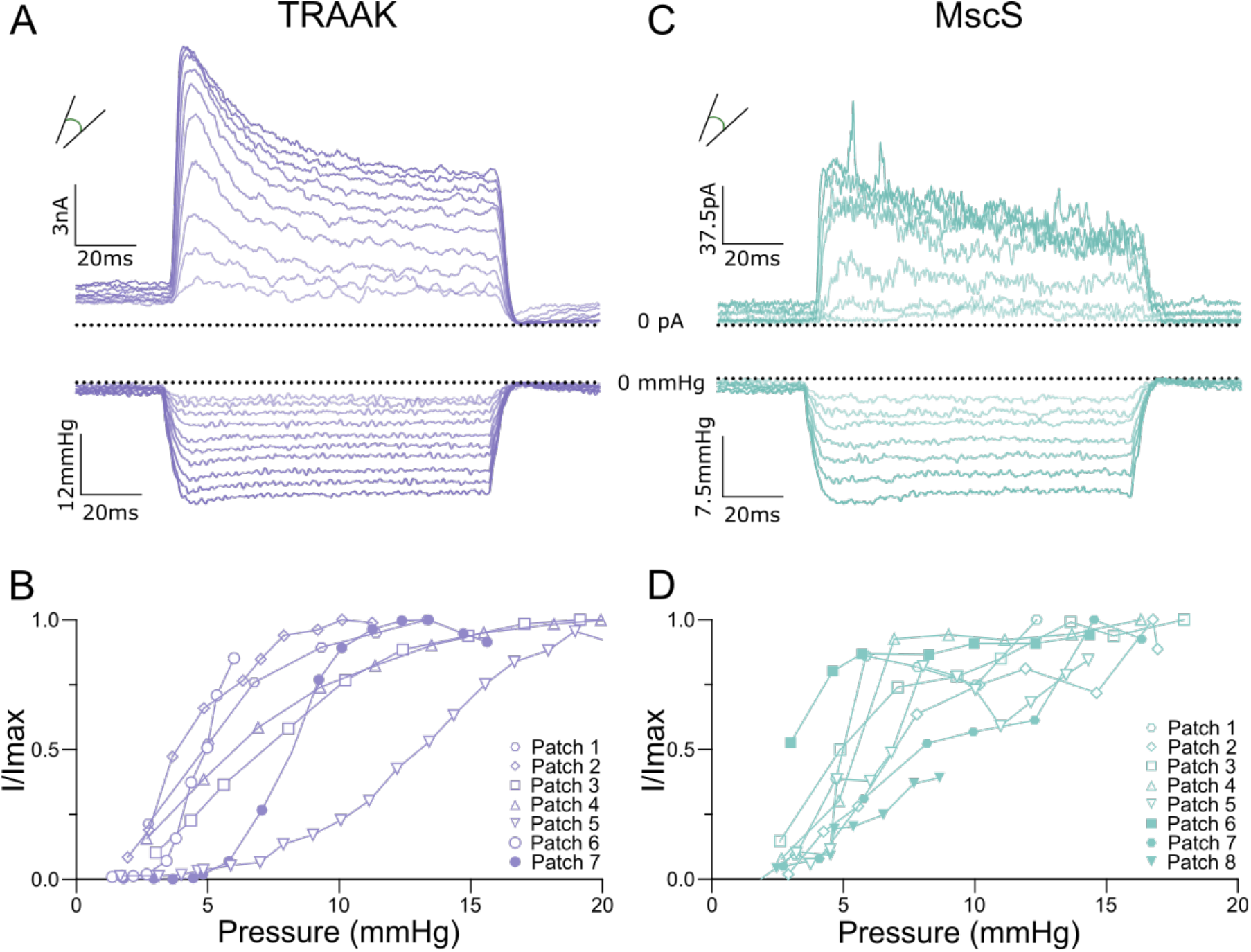
Pressure stimulation activates TRAAK and MscS channels. A. Macroscopic currents (upper) from a TRAAK-containing patch in response to a pressure step protocol (lower, V_hold_ = 0 mV, P_test_ = 0 to -25 mmHg, ΔP = -2.5 mmHg). B. Normalized current-pressure relationships from 7 TRAAK-containing patches. C. Macroscopic currents (upper) from a MscS-containing patch in response to a pressure step protocol (lower, V_hold_ = -60 mV, P_test_ = 0 to -14 mmHg, ΔP = -2 mmHg). D. Normalized current-pressure relationships from 8 MscS-containing patches.

Membrane tension (*T*) is related to the pressure difference across the patch (ΔP) and the membrane radius of curvature (r) according to the Young-Laplace equation: *T* = ΔPr/2. To calculate tension during mechanical stimulation, we visualized the membrane during pressure steps and measured the radius of patch curvature. Channels were co-expressed in oocytes with plasma membrane-targeted EGFP (EGFP fused to the CAAX lipidation motif from H-Ras). EGFP fluorescence was imaged during current recordings at a frame rate of 120 Hz. Pressure-induced changes in membrane curvature that were readily visible in TRAAK and MscS patches, with the magnitude of curvature change increasing with increasing pressure (Fig. 2A, Fig. S1). Patch radius of curvature was determined using a script input with manually identified points on the membrane (see Methods for details). Movie frames corresponding to the peak current elicited during each pressure step were analyzed.

**Figure 2.**
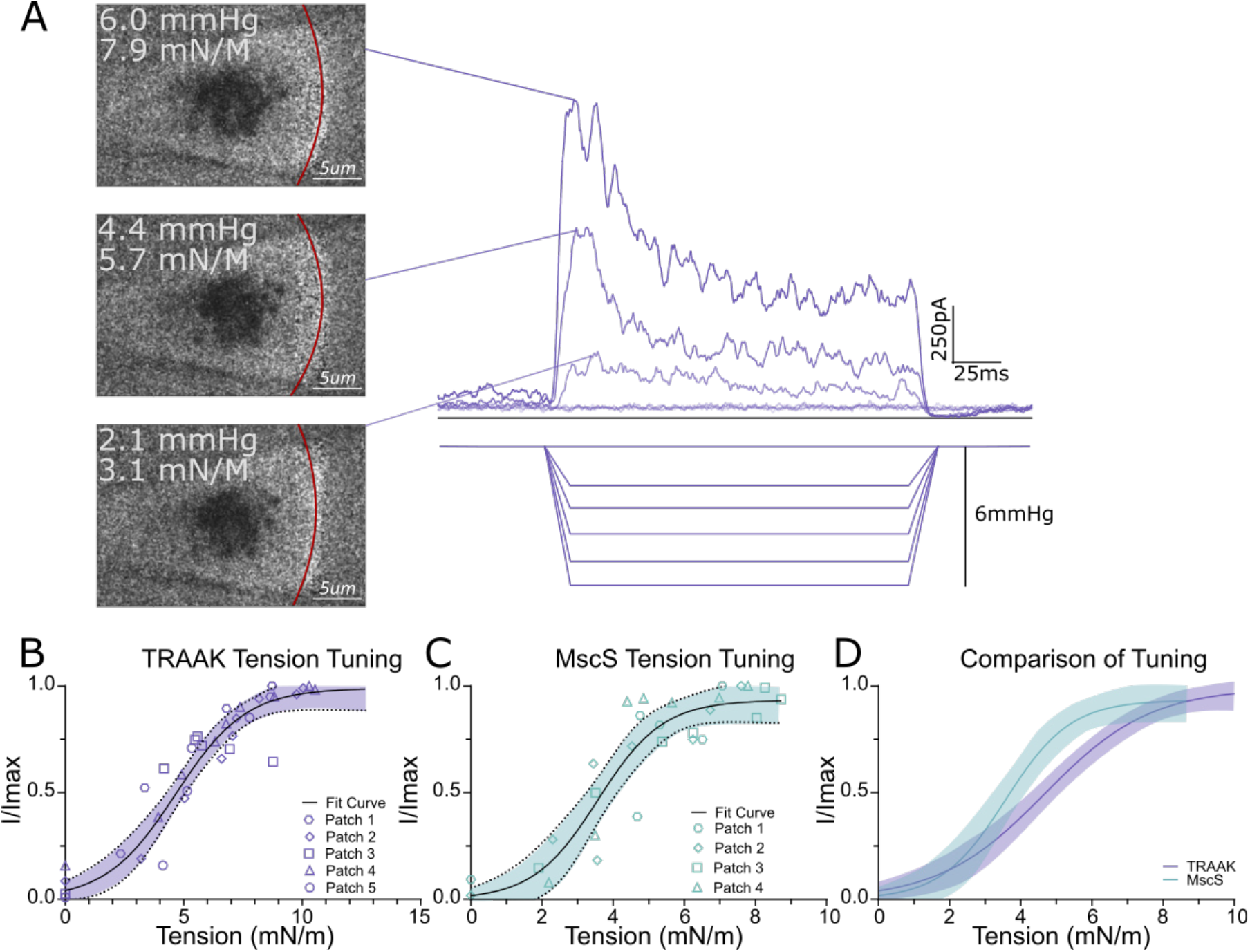
Quantification of TRAAK channel activation by membrane tension. A. (Right) Macroscopic currents from a TRAAK-containing patch in response to a pressure step protocol (V_hold_ = 0 mV, P_test_ = 0 to -6 mmHg, ΔP = -0.5 mmHg, 1 mmHg steps displayed). (Left) Fluorescent images of the patched membrane at the time of maximum current response during each pressure step. Fits for patch radius (red line), measured pressures, and calculated tensions are shown in each image. B. Normalized current-tension relationships for 5 TRAAK-containing patches. Global fit to a Boltzmann sigmoidal with 95% confidence intervals is shown. (T_50_ = 4.7 ± 0.2 mN/m, n = 5 patches). C. Normalized current-tension relationships for 4 MscS-containing patches. Global fit to a Boltzmann sigmoidal with 95% confidence intervals is shown (T_50_ = 3.7 mN/m ± 0.2 mN/m, n = 4 patches). D. Overlaid fits comparing TRAAK and MscS response to tension.

The tension response of TRAAK and MscS was consistent across patches (Fig. 2B, C, fit R^2^=0.87 and 0.83, respectively). TRAAK was activated by membrane tension over a broad range from low to near lytic tension. Data across patches were well fit to a Boltzmann with a midpoint *T*_50_=4.7 ± 0.2 mN/m and 10-90% range of 1.4-8.1 mN/m (mean ± SEM, n = 5 patches). TRAAK activity was indistinguishable when similar membrane tension was generated by positive or negative pressure stimulation that resulted in opposite membrane curvature (Fig. S1). MscS was activated with a midpoint *T*_50_=3.7 ± 0.2 mN/m and 10-90% of 1.2-6.3 mN/m (mean ± SEM, n = 4 patches). These results for MscS are consistent with values reported by other groups^9^, demonstrating the reliability of our approach. Comparing the tension response of TRAAK and MscS shows that the channels share a similarly low threshold for activation (defined by the 10% activation tension from Boltzman fits). However, TRAAK responds over a broader range of tension and has a higher *T*_50_ (Fig. 2D). In other words, the MscS response is steep and switch-like while the TRAAK response is more graded.

We next assessed ultrasound stimulation of TRAAK and MscS using the same patch imaging and recording setup. We designed a 3D-printed recording chamber to isolate the mechanical effects of ultrasound and ensure consistent ultrasound intensity at the patch (Fig. 3A). A 3.5 MHz ultrasound transducer was connected to the chamber through a mylar partition, and patched membranes were positioned at the point of maximum ultrasonic intensity. Channels were stimulated with 200 ms ultrasound bursts of increasing power from 0.04 W/cm^2^ to 5.4 W/cm^2^. As in our previous work^17^, we designed stimulation protocols to minimize bath temperature increases (to less than an estimated 0.05 ºC) to exclude potential thermal activation of channels.

**Figure 3.**
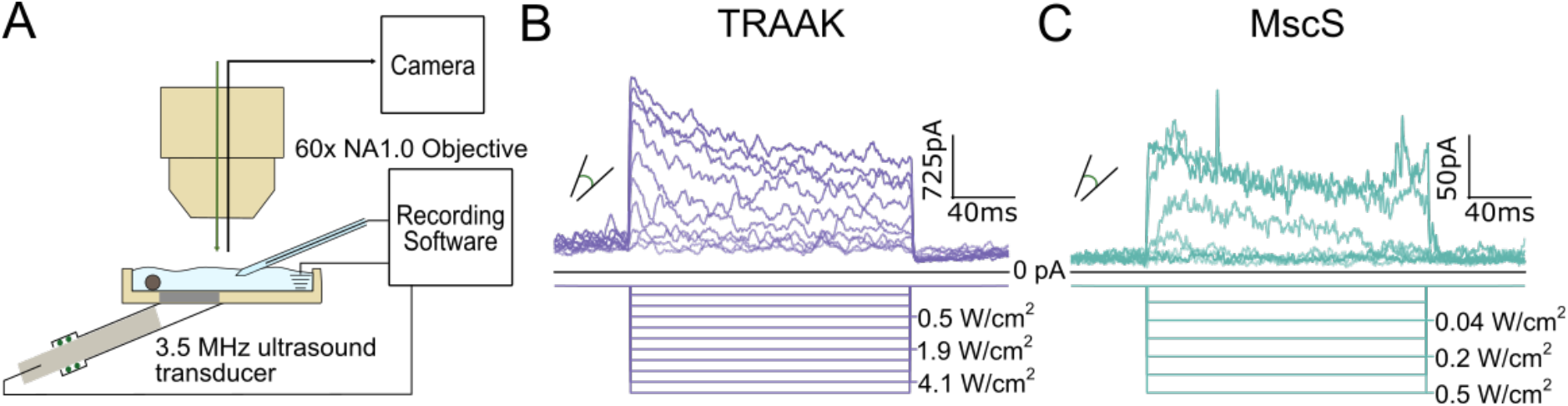
Ultrasound stimulation activates TRAAK and MscS channels. A. Schematic of recording setup for ultrasound stimulation during simultaneous patch recording and imaging. A 3.5 MHz ultrasound transducer is mounted into a custom 3D printed recording dish and patched membranes are positioned at the position of maximum ultrasonic power. B. Macroscopic currents from a TRAAK-containing patch in response to an ultrasound step protocol (V_hold_ = 0 mV, transducer driving voltage = 0-1 V, Δ driving voltage = 100mV, 100mV intervals displayed with measured power in W/cm^2^ at patch position indicated). C. Macroscopic currents from a MscS-containing patch in response to an ultrasound step protocol (V_hold_ = 0 mV, transducer driving voltage = 0-0.3 V, Δ driving voltage = 50 mV, measured power in W/cm^2^ at patch position indicated).

Increasing steps of ultrasound power increasingly activated TRAAK current (Fig. 3B) with a midpoint power of 2.2 ± 0.1 W/cm^2^ and a 10-90% range of 0.05-4.4 W/cm^2^ (mean ± SEM, n = 18 patches). At the highest ultrasound intensities tested, TRAAK was activated 42.6 ± 10.7-fold (mean ± SEM, n = 18 patches). MscS was similarly activated by ultrasound with a midpoint power of 1.0 ± 0.1 W/cm^2^ and a 10 to 90% range of 0.05-2.4 W/cm^2^ (mean ± SEM, n = 10 patches). At the highest ultrasound intensities tested, MscS was activated 22.9 ± 8.2-fold. The channels’ ultrasound responses mirror their tension responses; TRAAK and MscS have similar low thresholds for activation, but TRAAK shows a broader response range and higher midpoint power for activation.

Patch imaging showed that ultrasound stimulation, like pressure, induced membrane curvature changes that were concomitant with channel activation (Fig. 4A). In most patches, the membrane bowed outwards towards the pipette tip during ultrasound stimulation as observed for positive pressure application. The similarity of stimulus-induced curvature changes suggests that both ultrasound and pressure increase membrane tension to activate mechanosensitive TRAAK and MscS channels. We reasoned that if this is the mechanism for ultrasonic channel activation, then the membrane curvature of a patch should be the same when channels are activated to the same degree by pressure or ultrasound. Indeed, patch radii were indistinguishable when TRAAK was comparably activated by either pressure or ultrasound stimulus (Figure 4A-C, n=9 paired records from 5 patches, p = 0.18, paired t-test). Analysis of MscS gave the same result (Figure S2, n=7 paired records from 5 patches, p= 0.95, paired t-test). These results are consistent with ultrasound increasing membrane tension to activate mechanosensitive channels like canonical mechanical stimuli.

**Figure 4.**
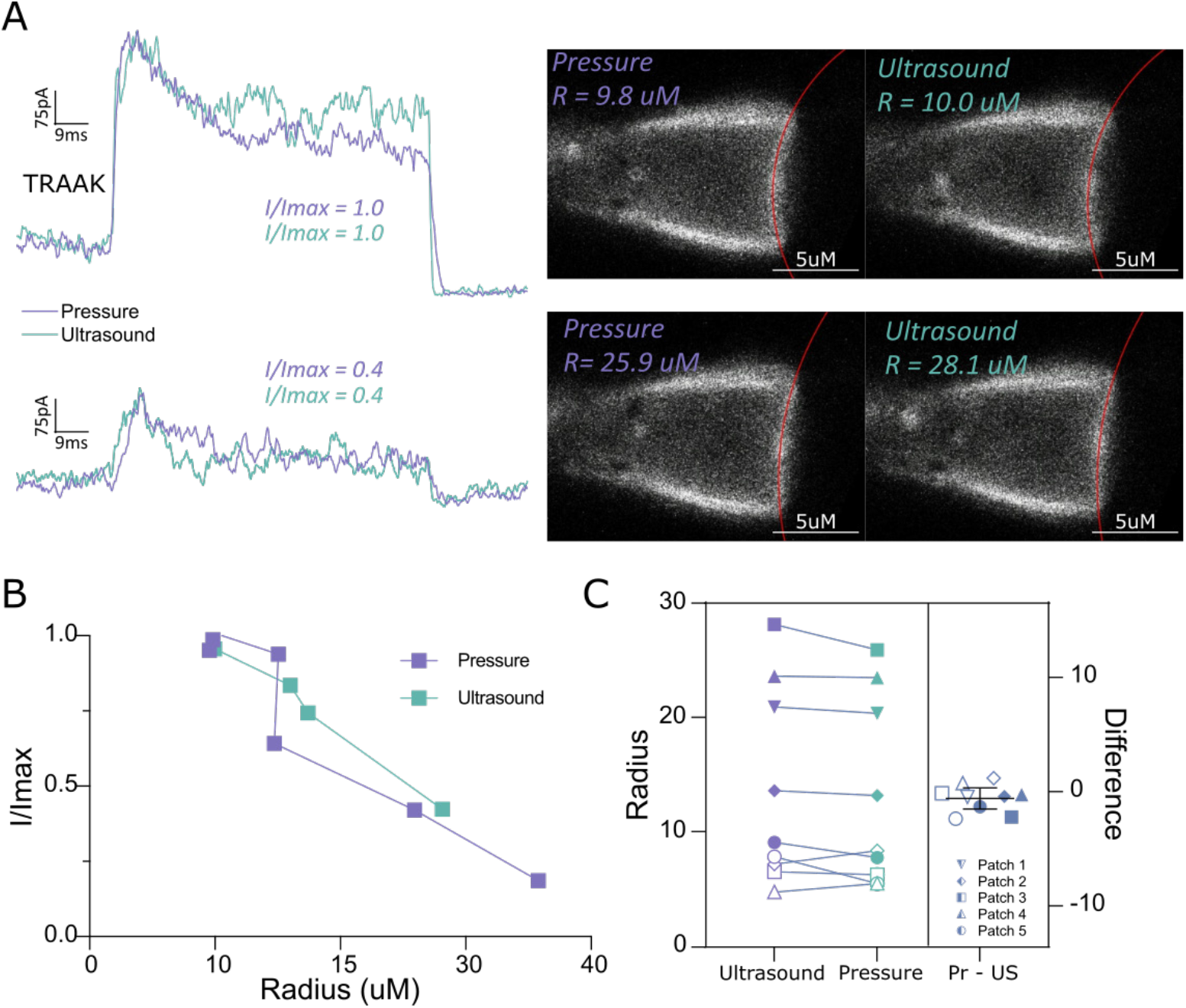
Ultrasound and pressure generate membrane tension to activate TRAAK channels. A. (left) Overlaid macroscopic currents and (right) corresponding fluorescent images from a TRAAK-containing patch in response to pressure (periwinkle) and ultrasound (green) stimulation. Traces and images are compared from the same patch activated to high (upper) and moderate (lower) I/Imax. Fits (red) and determined membrane radii are indicated on the images. Similar patch radii are observed when ultrasound or pressure stimuli generate similar channel activation. B. Normalized current-patch radius relationship calculated from a single patch in response to ultrasound and pressure stimulation. C. Comparison of patch radii observed during ultrasound or pressure stimulation that resulted in similar I/I_max_ values. Data from 5 different patches is shown (p = 0.18, paired t-test, not significant).

## Discussion

In this study, we used simultaneous patch imaging and recording to quantify the tension response of the mechanosensitive K^+^ channel TRAAK for the first time. We note that there are several caveats intrinsic to characterizing tension sensitivity in excised patches. First, we assume uniform tension across the patched membrane. Second, tension calculation requires hemispherical membranes (at infinite curvature, the Laplace-Young equation is undefined); it becomes increasingly challenging to calculate low tension values as patch radii of curvature increase in flatter membranes^32^. Third, membranes can have significant and variable basal tensions due to lipid adhesion to glass, estimated to be ∼0.5-4 mN/m in one study^33^. We attempted to minimize variability by pulling large diameter patches because they more frequently showed low basal curvature and low resting channel activity consistent with low basal tension^16,17^. Still, some variability in resting tension is likely to be present between patches and this is unaccounted for in our analysis. Large diameter patches also facilitated patch imaging and analysis, but this reduced the range of pressures that could be reached prior to patch rupture compared to prior work from our group and others^9,17^. This did not appear to bias results as similar tension responses were seen for MscS here and in previous work^9^.

We confirm TRAAK, like MscS, MscL, and Piezo1, is primarily a membrane tension-gated channel. TRAAK is activated by different pressures and membrane curvatures, including positive or negative pressures that curve the patch in opposite directions, but its tension response profile across patches is consistent. This supports the force-from-lipid model of mechanosensitivity in which open channel conformations are energetically favored in the presence of tension^3-4^. We can now compare the tension response of a mechanosensitive K2P channel to other channels. *E. coli* MscS has a low *T*_50_ of ∼2.5-7 mN/m measured from reconstituted proteoliposomes (depending on lipid composition)^5–9^. *E. coli MscL* has a high *T*_50_ of 11-12 mN/m measured from proteoliposomes or cell membranes^10–12^. Both bacterial channels show a steep tension response over a narrow tension range. The step-like opening is likely to be essential for the protective role of MscS and MscL as pressure-release valves in the bacterial response to osmotic shock^3,30^. Piezo1 is as or more sensitive than MscS with a *T*_50_ of ∼1.5-3 mN/m or ∼5 mN/m measured in on-cell patches or excised cell patches, respectively^12–14^. Piezo1, like MscS and MscL, responds over a narrow range of tension with a step-like response and may therefore serve as a switch to depolarize cells upon membrane stretch in numerous physiological contexts^1,2^. TRAAK is similarly sensitive to Piezo1 and MscS, with a threshold tension of ∼1 mN/m, but shows a notably broader response and correspondingly higher *T*_50_ of ∼5 mN/m. TRAAK is therefore expected to generate a K^+^ conductance proportional in magnitude to mechanical force across nearly the entire range of biologically feasible tension. Within a node of Ranvier, tension modulation of TRAAK activity could impact axonal excitability through, for example, altered repolarization rate, resting potential, or input resistance.

In the presence of membrane tension, expansion of protein cross-sectional area (ΔA) is favored by an energy equal to -*T*ΔA. If we assume area expansion is the dominant term driving opening of a mechanosensitive channel, ΔA and the intrinsic energy difference (ΔG) between an open and closed state can be derived from fitting tension response to the Boltzmann equation P_O_ = 1/1+exp((ΔG-*T*ΔA)/k_B_T)^32,34^. For TRAAK, we calculate a ΔA = 2.9 nm^2^ and ΔG = 13.49 × 10^−21^ J (∼3.3 k_B_T). Notably, ΔA derived from electrophysiological recordings is nearly identical to ΔA derived from experimental structures of open and closed TRAAK (ΔA = 2.7 nm^2^)^16,18^. This provides further support for the structural model in which mechanical opening involves movement of TRAAK transmembrane helix 4 to an “up” conformation, expanding channel cross-sectional area and sealing membrane-facing openings to lipid block^16,18^.

We also utilized the patch imaging and recording setup developed here to gain insight into the basis of ultrasound activation of mechanosensitive channels. Among neuromodulatory techniques, ultrasound is uniquely penetrant and focusable in biological tissues. This means ultrasound can, for example, be targeted non-invasively to deep-brain structures through the skull with excitatory or inhibitory effects on neural activity that depend on the target and stimulus parameters^25,26^. Recent studies have implicated mechanosensitive channels as mediators of some ultrasound effects with channel activation observed in vitro, upon heterologous expression, and from endogenously-expressed channels in central and peripheral neurons^17,27–29,35–37^. Harnessing and predicting ultrasound activation requires a complete understanding of the underlying molecular mechanisms involved.

We demonstrated both TRAAK and MscS channels are activated by ultrasound in excised patches. We observed similar response profiles to ultrasound and pressure stimulation for each channel and found that both stimuli generate changes in patch curvature correlated with channel activation. Within a single patch, ultrasound and pressure stimulation are indistinguishable in generating a relationship between membrane radius of curvature and channel activity, suggesting both stimuli activate channels through membrane tension. This is consistent with our previous work showing ultrasound and pressure promote the same mechanically gated TRAAK open state, distinguished from the alternative leak open state by a higher conductance and longer duration^17^. Together, these results suggest the principal mechanism by which ultrasound activates mechanosensitive channels is by increasing membrane tension. Other potential effects, including temperature increase, cavitation, or acoustic scattering, may be relevant under other conditions^25,26^, but are not likely to explain channel activation in our experiments. Instead, acoustic radiation force and resulting acoustic streaming likely account for mechanical activation of channels by ultrasound^25,35,38^. Indeed, work in other systems has demonstrated analogous mechanical displacements of reconstituted lipid bilayers and cell membranes^38,39^.

Recent work has used a range of ultrasound stimulation parameters to activate mechanosensitive channels with varying frequencies (from 300 kHz to 42 MHz), powers (0.05 to 750 w/cm^2^), beam profiles (with focal points of 0.2 to 4 mm^3^), and waveforms (e.g., durations from 10-200 ms), producing varying effects^17,26–29,35–37^. We note a substantial difference in midpoint power for TRAAK activation in this work (2.2 W/cm^2^, 3.5 MHz) and our prior study (0.8 W/cm^2^, 5 MHz)^17^, which may be explained by the difference in stimulation frequency. Screening stimulation protocols to maximize the mechanical effects of ultrasound could therefore provide further insight into underlying physical mechanisms of TRAAK activation and be useful for minimizing off-target effects in neuromodulatory and sonogenetic applications.

## Methods

### Expression in and recording from *Xenopus laevis* oocytes

A construct encoding full-length *Homo sapiens* TRAAK (UniProt Q9NYG8-1) was codon optimized to enhance eukaryotic expression in *P. pastoris, S. frugiperda*, and *H. sapiens* (without changing the native amino acid sequence), synthesized (Genewiz), and cloned into a modified pGEMHE vector using Xho1 and EcoR1 restriction sites. The transcribed message encodes *H. sapiens* TRAAK amino acids 1 to 393 with an additional amino acid sequence of “SNS” at the C terminus. The coding sequence for *E. coli* MscS^31,40^ was cloned into a modified pGEMHE vector using EcoR1 and Xho1 restriction sites. The transcribed message encodes pFLAG-CTC-GFP-MscS^40^ amino acids 1-286 with an attached GFP upstream of the MscS ATG. A construct encoding EGFP fused to the CAAX-containing C-terminal tail of *H. sapiens* HRas (NP_005334 amino acids 170-189) through a GGRS linker was cloned into a pCS2+ vector with a Kozak sequence of CACC using Gibson assembly. Complementary DNA (cDNA) was transcribed from these plasmids *in vitro* using T7 polymerase and 0.1 to 10 ng (TRAAK and MscS) and 3-10 ng (EGFP-CAAX) complementary RNA (cRNA) in 50 nL H_2_O was injected into *Xenopus laevis* oocytes extracted from anesthetized frogs. Currents were recorded at 25 °C from inside-out patches excised from oocytes 1 to 5 d after cRNA injection. The pipette solution contained the following: 15 mM KCl, 135 mM NaCl, 2 mM MgCl2, 10 mM HEPES, pH = 7.4 with KOH and the bath solution contained 150 mM KCl, 2 mM MgCl2, 10 mM HEPES, 1 mM EGTA, pH = 7.1 with KOH. Currents were recorded using an Axopatch 200B Patch Clamp amplifier at a bandwidth of 1 kHz and digitized with an Axon Digidata 1550B at 500 kHz. Pressure was applied with a second-generation high-speed pressure clamp device (HSPC-2-SB, ALA Scientific Instruments).

### Ultrasound setup and application

We conducted both pressure and ultrasound recordings in a custom 3D printed mount that placed the ultrasound transducer in line with the patch pipette at a 20° angle. The recording chamber, containing the ultrasound transducer mount and recording bath, was made from a clear SLA photopolymer (Formlabs RS-F2-GPCL-04). To ensure the bath fluid would not leak out of the chamber, the transducer mount was fitted with two nitrile O-rings (SUR&R 55AA89). Inside-out patches were excised from oocytes within the ultrasound chamber. The patch was centrally positioned ∼1 in (25.4 mm) away from the cylindrical transducer base surface, separated by bath solution and a thin sheet of mylar which the oocyte rested upon. An ultrasound wave was generated using a V326-SU (Olympus) focused-immersion ultrasonic transducer with a 0.375 in (9.525 mm) nominal element diameter, which had a focal point at 25.2 mm (0.993 in) and an output center frequency of 3.5 MHz. To trigger the transducer’s ultrasound pulses, a function generator (Agilent Technologies, model 33220A) was used to send an input voltage waveform to an ENI RF (radio frequency) amplifier (model 403LA), which provided the output power to the ultrasound transducer for producing the acoustic pressure profile of a stimulus waveform. The timing of the ultrasound stimuli was controlled by triggering the function generator manually or by software (Clampex 10.7). In the case of ultrasound pulse generation though software, a Clampex 10.7-generated waveform triggered a first function generator through a digitizer (Axon Digidata 1550B), which triggered a second function generator, which triggered the RF amplifier that drives the ultrasound transducer. Solutions were degassed to minimize microbubble cavitation and ultrasound attenuation.

### Calculating ultrasound pressure and power

The output pressures were measured using a calibrated hydrophone (Onda, model HNR-0500). The hydrophone measurements were performed at the position of peak spatial pressure. When converting the measured voltages into pressures, we accounted for the hydrophone capacitance according to the manufacturer’s calibration. Using the appropriate conversion factor listed under the Pascals-per-volt column on the look-up table that was supplied with the calibrated hydrophone, the hydrophone voltage-trace waveform was transformed into an acoustic-pressure waveform measured in MPa. We calculated the ultrasound power intensity in Watts/square centimeter (W/cm^2^) with the following equation:

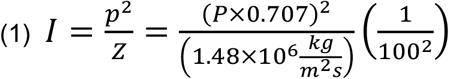

### Patch imaging and membrane tension calculation

Excised patches were illuminated with a white LED through a GFP filter and water immersion objective lens (x60, NA1.0). Movies were recorded at 120 Hz with an infrared camera (IR-2000, DAGE-MTI). Images were analyzed within FIJI (ImageJ). Image contrast was enhanced to facilitate analysis. Video files were loaded into FIJI and converted into a JPEG stack. Frames were then time-matched to stimuli by multiplying frame rate and time. Frames of interest were selected, three points were chosen along the curve of the membrane, and the coordinates of all three points were loaded into a python script utilizing the following equations:

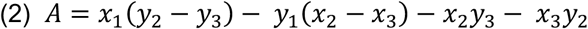

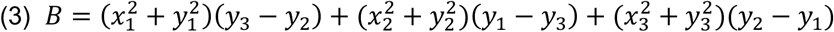

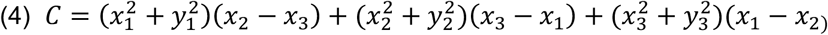

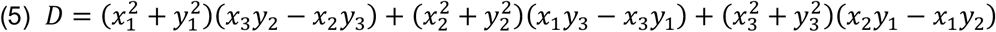

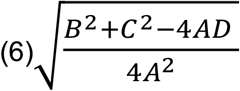

Radii were used to calculate membrane tension using Laplace’s law as previously described^7,8,12^.

## Author contributions

B.S., T.A.D., and V.P. performed electrophysiology. R.A.R. contributed to early stages of the project. T.A.D. and R.A.R. generated constructs and wrote scripts for data analysis. B.S. and T.A.D. analyzed data. T.A.D. generated figures. T.A.D. and S.G.B. wrote the paper with input from all authors. S.G.B. supervised the project.

## Acknowledgements

We thank Dr. E. Iscaoff, Dr. Y. Yu, A. Chou, and C. Stanley for providing *Xenopus* oocytes; Dr. E. Haswell for the MscS construct; Dr. E. Iscaoff and Dr. A. Winans for the GFP-CAAX construct; and members of the Brohawn laboratory for critical review and discussions, especially K. Tucker, Dr. D. Kern, and Dr. A. Elleman. S.G.B. is a New York Stem Cell Foundation-Robertson Neuroscience Investigators. This work was supported by the New York Stem Cell Foundation (S.G.B.) and NIH/NINDS K99 NS125102 (B.S.).

---

**Supplemental Figure 1.**
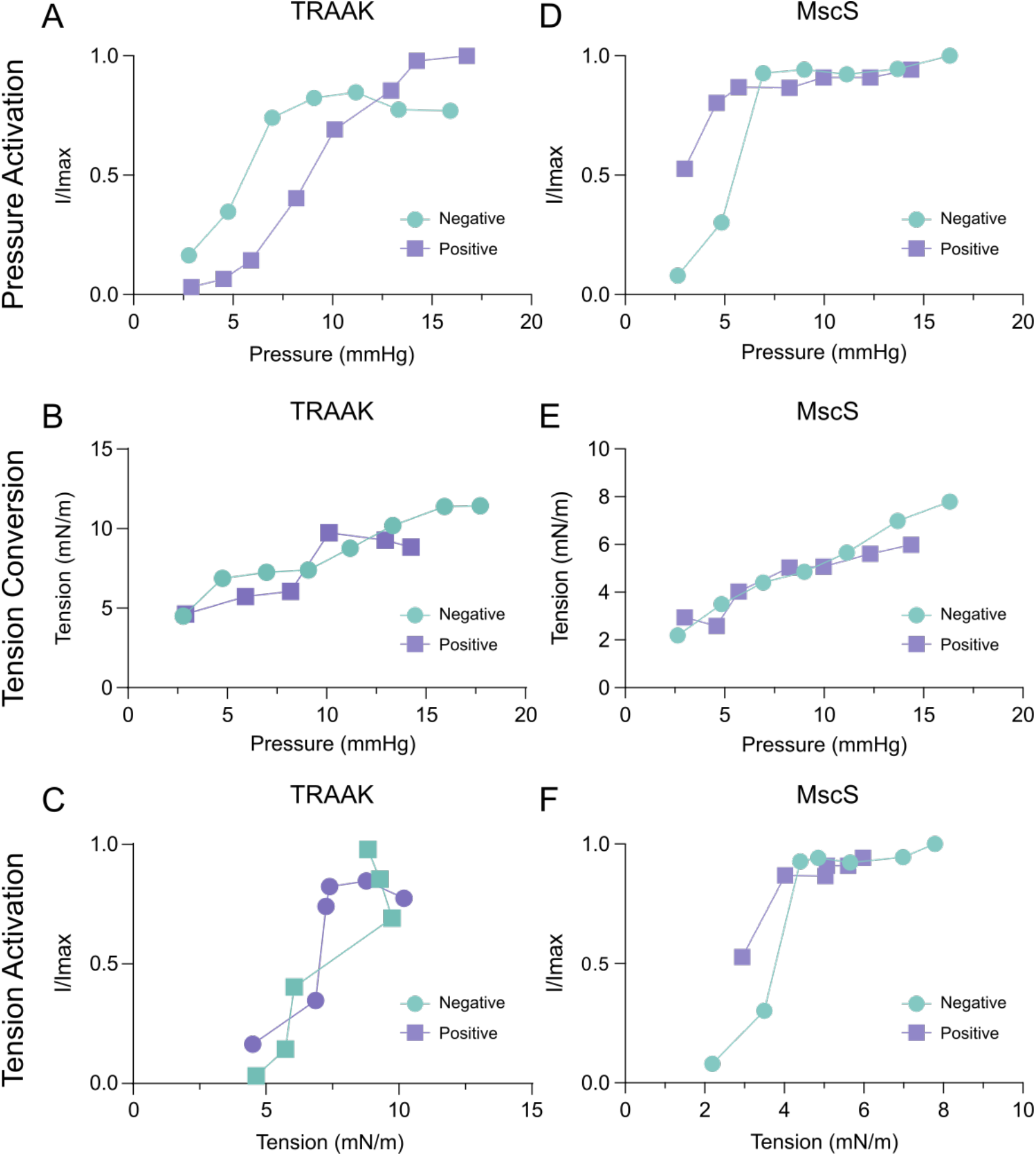
TRAAK and MscS are equivalently activated by tension generated by positive and negative pressure. A. Normalized current-pressure relationship for positive (periwinkle) and negative (green) pressure for a TRAAK-containing patch. B. Tension-pressure relationship from data in A. C. Normalized current-tension relationship for channels activated by positive (periwinkle) and negative (green) pressure. D-F. Same as A-B, but for a MscS-containing patch.

**Supplemental Figure 2.**
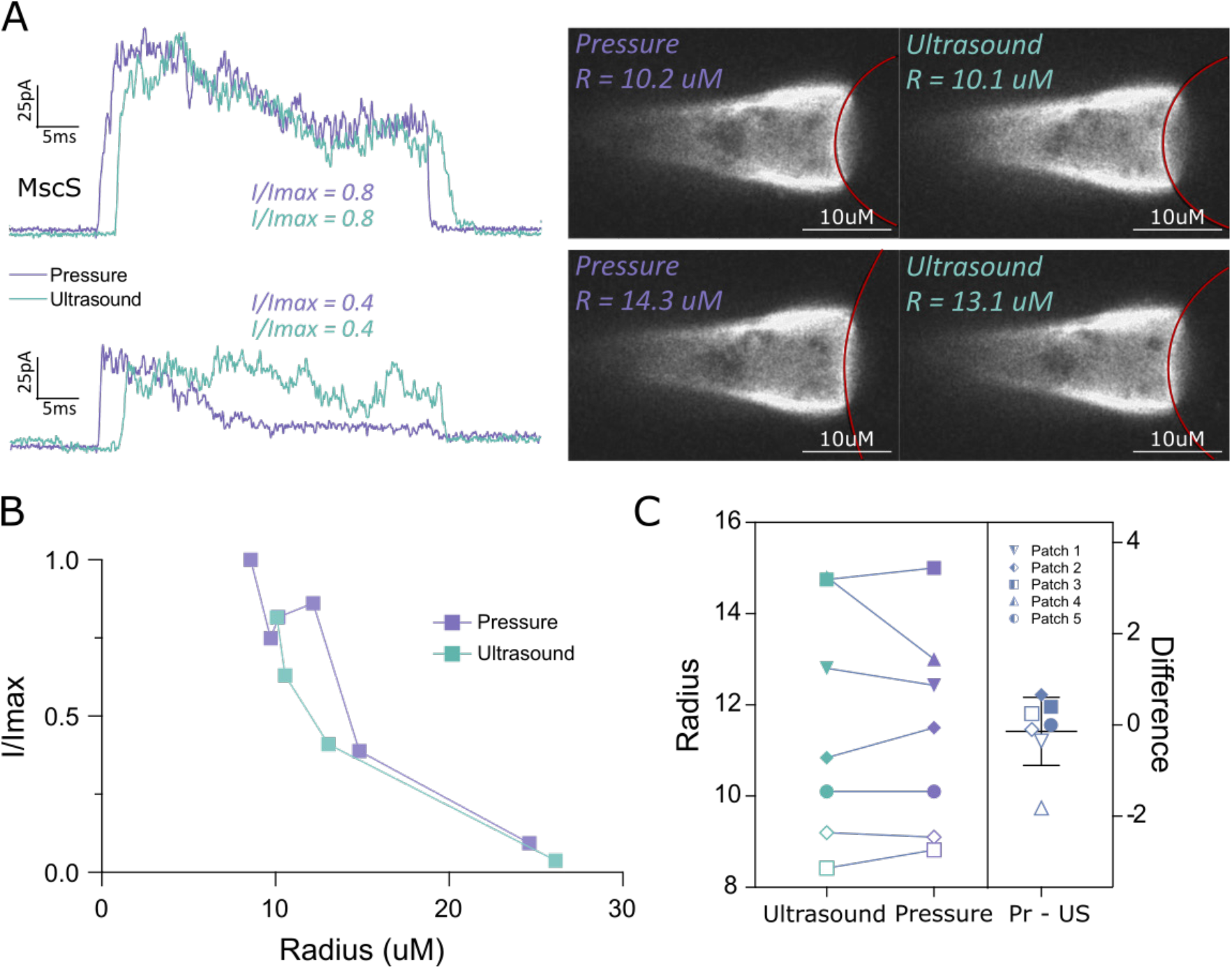
Ultrasound and pressure generate membrane tension to activate MscS channels. A. (left) Overlaid macroscopic currents and (right) corresponding fluorescent images from a MscS-containing patch in response to pressure (periwinkle) and ultrasound (green) stimulation. Traces and images are compared from the same patch activated to high (upper) and moderate (lower) I/Imax. Fits (red) and determined membrane radii are indicated on the images. Similar patch radii are observed when ultrasound or pressure stimuli generate similar channel activation. B. Normalized current-patch radius relationship calculated from a single patch in response ultrasound and pressure stimulation. C. Comparison of patch radii observed during ultrasound or pressure stimulation that resulted in I/I_max_ values near 0.5. Data from 5 different patches is shown (p = 0.95, paired t-test, not significant).

